# Bidirectional multiciliated cell extrusion is controlled by Notch driven basal extrusion and Piezo 1 driven apical extrusion

**DOI:** 10.1101/2023.01.12.523838

**Authors:** Rosa Ventrella, Sun K. Kim, Jennifer Sheridan, Aline Grata, Enzo Bresteau, Osama Hassan, Eve E. Suva, Peter Walentek, Brian Mitchell

**Affiliations:** Northwestern University, Feinberg School of Medicine, Department of Cell and Developmental Biology; Current position; Assistant professor, Precision Medicine Program, Midwestern University; University of Freiburg, Renal Division, Internal Medicine IV, Medical Center and CIBSS Centre for Integrative Biological Signalling Studies; Northwestern University, Lurie Cancer Center

**Keywords:** Multiciliated cells, cell extrusion, Notch, Piezo 1

## Abstract

*Xenopus* embryos are covered with a complex epithelium containing numerous multiciliated cells (MCCs). During late stage development there is a dramatic remodeling of the epithelium that involves the complete loss of MCCs. Cell extrusion is a well-characterized process for driving cell loss while maintaining epithelial barrier function. Normal cell extrusion is typically unidirectional whereas bidirectional extrusion is often associated with disease (e.g. cancer). We describe two distinct mechanisms for MCC extrusion, a basal extrusion driven by Notch signaling and an apical extrusion driven by Piezo1. Early in the process there is a strong bias towards basal extrusion, but as development continues there is a shift towards apical extrusion. Importantly, receptivity to the Notch signal is age-dependent and governed by the maintenance of the MCC transcriptional program such that extension of this program is protective against cell loss. In contrast, later apical extrusion is regulated by Piezo 1 such that premature activation of Piezo 1 leads to early extrusion while blocking Piezo 1 leads to MCC maintenance. Distinct mechansms for MCC loss underlie the importance of their removal during epithelial remodeling.

**Summay Statement:** Cell extrusion typically occurs unidirectionally. We have identified a single population of multiciliated cells that extrudes bidirectionally: Notch-driven basal extrusion and Piezo 1-mediated apical extrusion.

## Introduction

Ciliated epithelial-driven fluid flow is essential for moving mucus and particles across the lungs and oviduct as well as for cerebrospinal fluid circulation. It is also required for the locomotion of aquatic animals. In many of these biological contexts, a ciliated epithelium must maintain homeostasis and undergo epithelial remodeling involving the loss and replenishing of multi-ciliated cells (MCCs) (Rock et al., 2011; Walentek, 2021). *Xenopus* embryos contain a ciliated epithelium comprising numerous MCCs that promote both locomotion and gas exchange prior to lung development. During their development, the need for cilia-driven flow is transient and is ultimately replaced by the swimming motion of the more mature tadpoles. This transition requires a dramatic remodeling of the epithelium that involves the complete loss of MCCs, while maintaining epithelial barrier function.

Cell extrusion is a well-characterized process in maintaining epithelial homeostasis and occurs during the remodeling of numerous tissues (Andrade and Rosenblatt, 2011; Fadul and Rosenblatt, 2018; Gudipaty et al., 2018; Ohsawa et al., 2018). Cell extrusion from an epithelium takes place in multiple forms, including the loss of unwanted apoptotic cells, the loss of less competitive cells, and the stochastic loss of cells due to overcrowding (Atieh et al., 2021; Eisenhoffer et al., 2012; Ellis et al., 2019; Fadul et al., 2021; Gu et al., 2011; Levayer et al., 2016; Slattum et al., 2014). This removal of cells can occur either basally or apically, depending on the context (Nanavati et al., 2020). While there is considerable variation across model systems and organ systems, it is common that for a particular tissue, cell extrusion typically occurs either apically or basally, but rarely both. Importantly, the incorrect orientation of cell extrusion is often associated with disease states and is well characterized in cancer and hyperplasia (Gu et al., 2015; Gudipaty and Rosenblatt, 2017). Previously, it has been shown that sphingosine 1-phosphate(S1P)-Rho mediated signaling is required for cell extrusion of both apoptotic and non-apoptotic cells (Eisenhoffer *et al*., 2012; Gudipaty and Rosenblatt, 2017). Additionally, the stretch-activated ion channel Piezo 1 has been shown to act as a mechanosensor of epithelial tissue remodeling (Gudipaty et al., 2017; Miyamoto et al., 2014; Peyronnet et al., 2013; Stewart and Davis, 2019) and has been identified as an upstream factor that drives live cell extrusion (Eisenhoffer *et al*., 2012).

The transcriptional regulators of the MCC lineage are well characterized (Boon et al., 2014; Collins et al., 2021b; Ma et al., 2014; Stubbs et al., 2012; Terre et al., 2016; Vladar and Mitchell, 2016). In particular, members of the Geminin family of proteins, GemC1 and MCIDAS, form complexes with DP1 and E2F4/5 to promote the transcriptional regulation of the MCC fate (Ma *et al*., 2014; Terre *et al*., 2016). In fact, in the proper context, GemC1 and MCIDAS are sufficient and necessary to drive the formation of MCCs (Boon *et al*., 2014; Kim et al., 2018; Ma *et al*., 2014; Stubbs *et al*., 2012; Terre *et al*., 2016). Interestingly, both GemC1 and MCIDAS are downregulated after the MCC fate is established, whereas downstream transcription factors that drive ciliogenesis, such as RFX2 and FoxJ1, continue to be expressed throughout the life of the MCC (Briggs et al., 2018; Chung et al., 2014; Chung et al., 2012; Quigley and Kintner, 2017; Stubbs *et al*., 2012).

While GemC1 and MCIDAS promote MCC specification, Notch signaling acts as a negative regulator of MCC formation. In mammalian lungs and *Xenopus* embryonic skin, activation of Notch signaling inhibits MCC formation, while inhibition of Notch results in increased MCC differentiation (Deblandre et al., 1999; Lewis and Stracker, 2021; Liu et al., 2007; Rock *et al*., 2011). Additionally, the loss of MCCs from the older *Xenopus* epithelium has been shown to require Notch signaling that has been proposed to emanate from the developing lateral line primordium and the underlying mesoderm based on the expression of Notch ligands (Tasca et al., 2021). Overexpression of the active Notch intracellular domain (NICD) in MCCs has been shown to increase the number of TUNEL staining positive cells, indicating that Notch can induce apoptosis of MCCs, leading to a loss of MCCs. Furthermore, the Notch signaling pathway in mouse has been proposed to regulate the transdifferentiation of club cells into MCCs in adult tissue in which Notch signaling has been blocked (Lafkas et al., 2015).

In this study, we show two distinct mechanisms of MCC extrusion during *Xenopus* embryonic skin remodeling; a Notch based basal extrusion that requires a cell-autonomous licensing event tied to the loss of MCC transcriptional commitment and a separate apical extrusion that is regulated by the mechanosensory channel Peizo 1. These distinct mechanisms work in concert to ensure that all MCCs are lost from the epithelium by ST48 of *Xenopus* embryonic development.

## Results

### Loss of fluid low and multiciliated cells by ST48

MCCs work together to drive a directed fluid flow across the surface of the epithelium during *Xenopus* embryonic development. To examine how long MCC-driven fluid flow is maintained, we measured the displacement of fluorescent beads across the embryo’s surface (Mitchell et al., 2007; Werner and Mitchell, 2013). We performed a developmental time course of this rate and found that the flow rate peaks 4 days post fertilization (dpf) at stage 38 (ST38). Then it steadily decreases over the next few days and is essentially gone by 9 dpf at ST48 (Figure 1A). This loss of flow corresponds with the progressive loss of cilia observed with acetylated tubulin antibody staining between ST38 and ST48, as previously reported (Figure 1B-C) (Tasca *et al*., 2021). Importantly, using a transgenic line of *Xenopus* that expresses membrane-bound RFP (memRFP) specifically in MCCs driven by the alpha-tubulin A1A (Tub) promoter (TgTub-memRFP), we see a similar time frame of MCC loss (Figure 1B-C) (Brooks and Wallingford, 2015; Collins et al., 2021a). We find that the RFP labeling of these cells is quite stable and we can trace MCCs even as they delaminate from the epithelium and undergo cell death (Figure 3A,C and Movie S1). The RFP-positive remnants of these cells continue to be visible after delamination and death. Long-term imaging using light-sheet microscopy shows cell remnants for longer than 24 HRs suggesting that the mem-RFP labeling is robust and stable. At later stages of development, we do occasionally see some RFP-positive cells devoid of acetylated tubulin, which is consistent with the previous claim that some MCCs are undergoing a trans- or de-differentiation event (Tasca *et al*., 2021). However, the complete loss of RFP positive cells by ST48 coupled with the prolonged presence of stable mem-RFP positive cell debris found in the epithelium suggests that this change of fate is transient and that ultimately all of the MCCs are lost from the epithelium.

**Figure 1:**
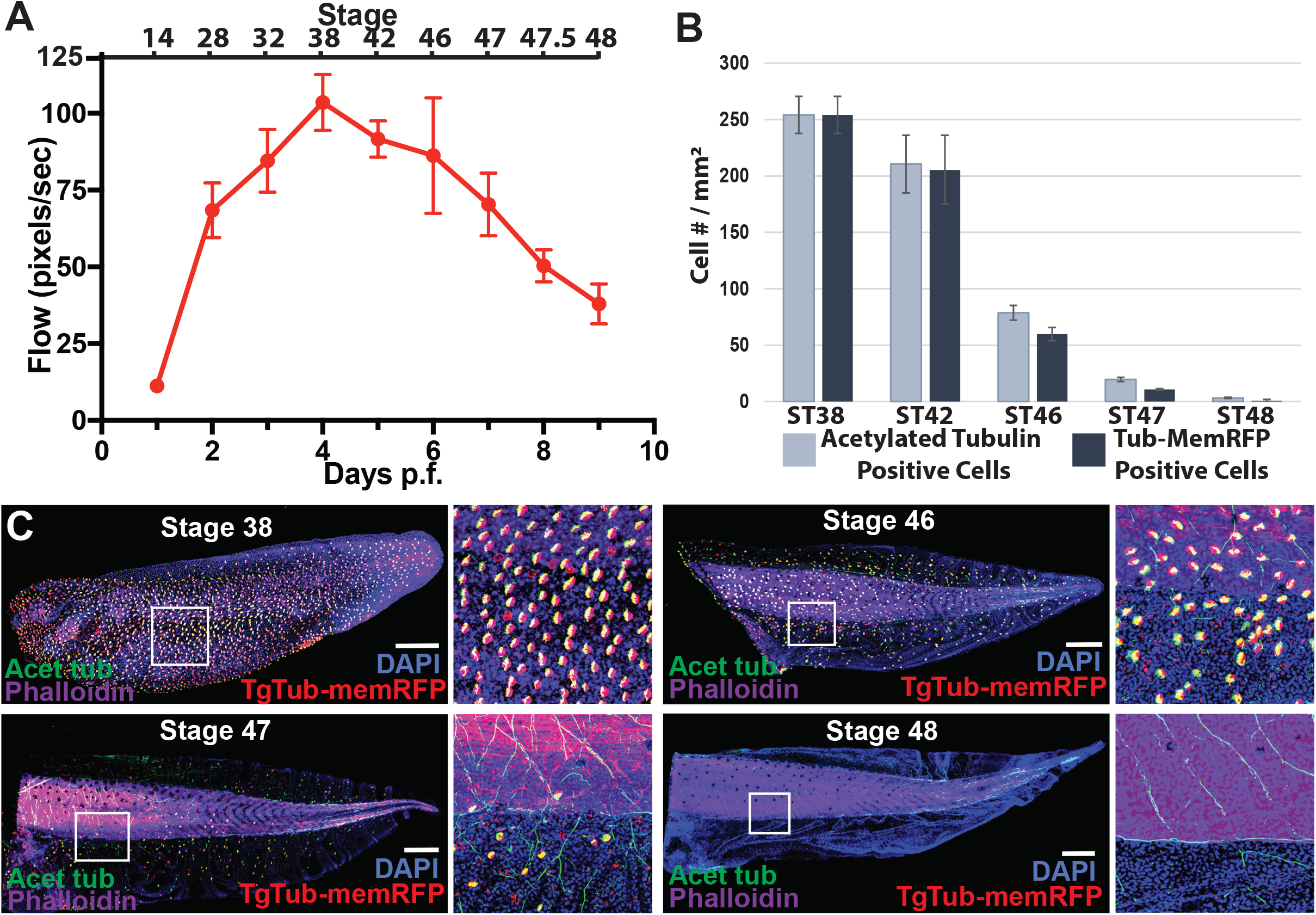
The loss of cilia driven fluid flow accompanies the loss of Multi-ciliated cells. (A) Quantification of cilia driven fluid flow as measured by fluorescent bead displacement across the surface of the epithelium showing that flow peaks at 4 dpf and is essentially lost by 9 dpf, n=8 animals / time point. (B) Quantification of MCC number using both the transgenic line TgTub-memRFP driving MCC specific expression of Membrane RFP and antibody staining of acetylated tubulin. (C) Representative images of the progression of MCC loss in embryos between ST38 and ST48 showing acetylated Tub (green), TgTub-memRFP (red), and phalloidin (purple) and DAPI (blue). Scale bar is 500μm.

**Figure 2:**
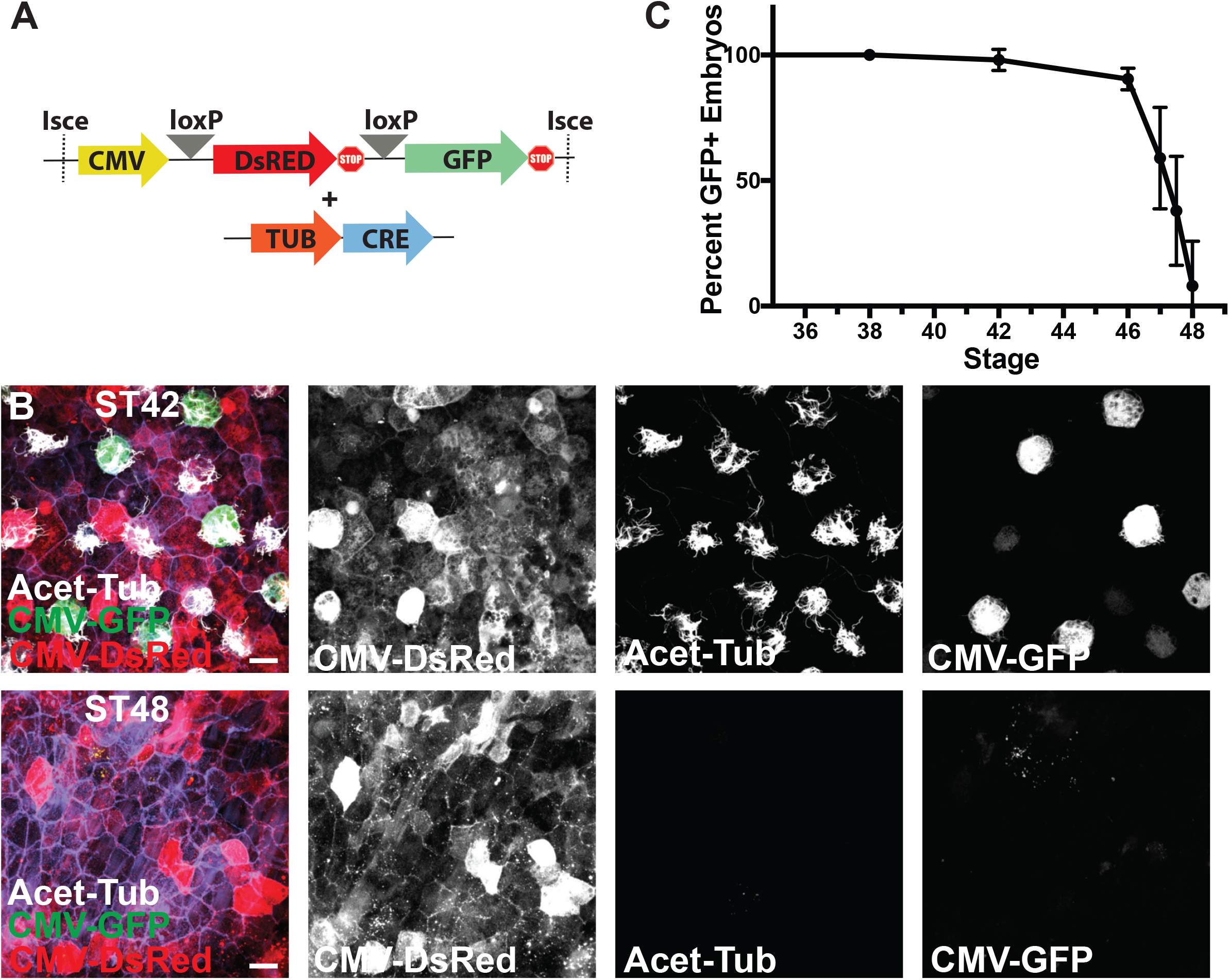
Lineage tracing of MCC fate. (A) Experimental design showing lineage tracing constructs of the CMV RFP loxP GFP cassette together with the Tub driven CRE. (B) Lineage tracing experiment showing the conversion of RFP to GFP in ST42 MCCs upon expression of CRE specifically in MCCs under control of the Tub promoter, and the loss of MCC specific CMV expression of GFP despite the maintenance of broad CMV driven RFP expression at ST48. (C) Developmental quantification of embryos containing GFP positive cells in lineage tracing experiments, n=102 trasngenic animals over 5 independent experiments. Scale bar is 20μm.

**Figure 3:**
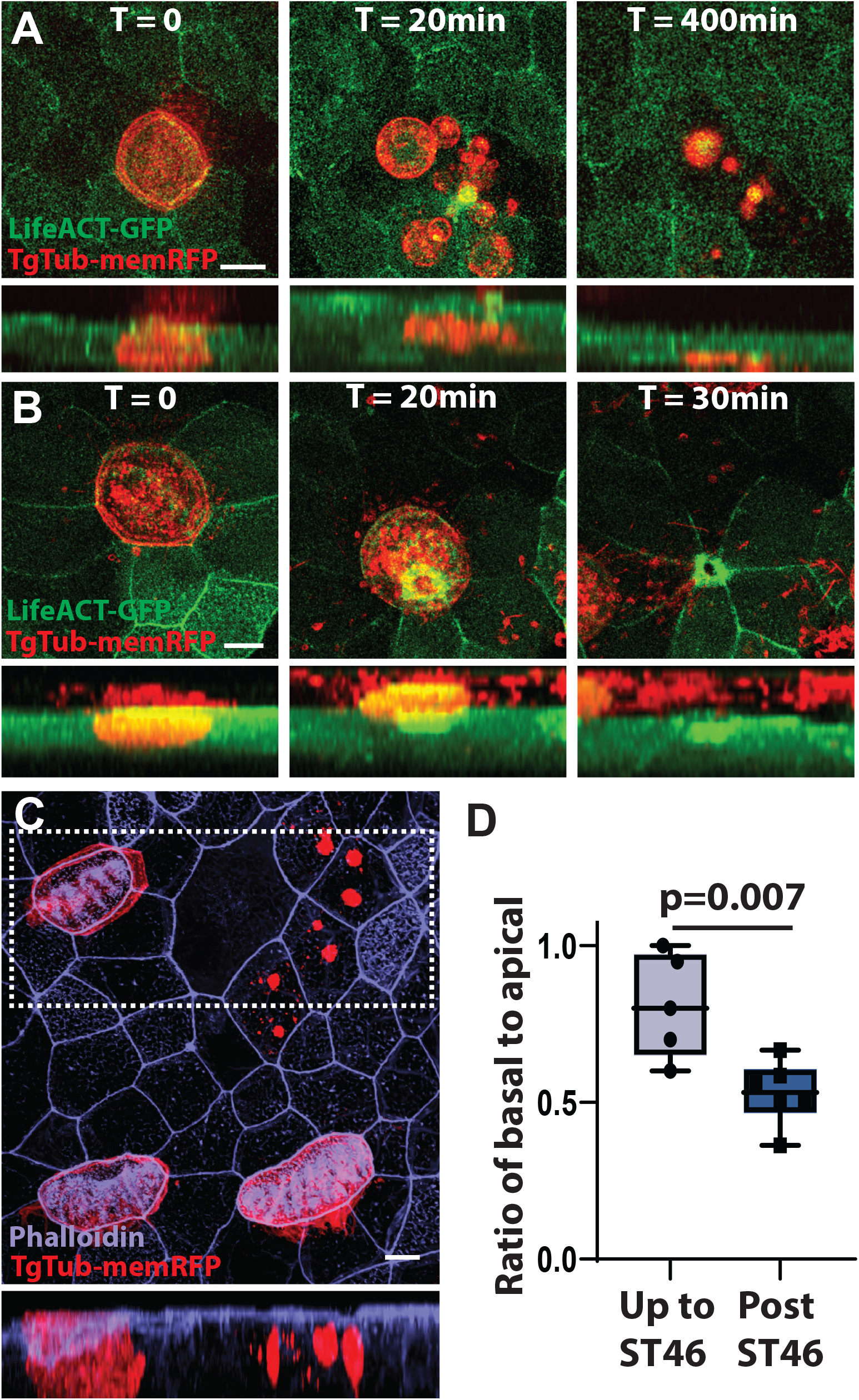
MCCs are extruded both basally and apically. (A) Representative example of a time lapse movie of TgTub-memRFP embryos injected with LifeACT-GFP showing an MCC undergoing basal extrusion with the RFP positive cell remnants remaining visable for at least 400 min. See Movie S1. (B) Representative example of a time lapse movie showing an MCC undergoing apical extrusion. See Movie S2 (C) Representative example of RFP positive MCC remnants in fixed tissue stained with phalloidin. (D) Quantification of MCC loss via long-term light-sheet imaging showing the ratio of basal to apical extrusion from ST42 to ST46 and after ST46, n=132 extrusion events from 5 embryos up to ST46, and 69 events from 6 embryos post ST46, p = 0.007. See Movie S3. Scale bars are 10μm.

### Genetic lineage tracing of MCC disappearance

While the RFP positive labeling of MCCs and the remnants of dead MCCs using the transgenic line was consistently robust, one could argue that as MCCs alter their fate, the tubulin promoter shuts off causing loss of mem-RFP expression from these cells, which would obscure our interpretation that all MCCs are lost. To rule out this possibility, we generated F0 transgenic embryos using a Cre-LoxP-based lineage tracing cassette in which a CMV promoter drives ubiquitous expression of DsRed with a stop codon, flanked by LoxP sites, and followed by GFP (Figure 2A) (Rankin et al., 2009; Werdien et al., 2001). In the absence of CRE, CMV drives the expression of DsRed throughout the epithelium without any detectible green (Werdien *et al*., 2001). In contrast, when we introduce CRE specifically in MCCs using the Tub promoter, we see a strong conversion of Red to Green, indicating DsRed excision. Importantly, the MCC specific GFP expression is still driven by the CMV promoter (Figure 2B). At ST42, we find many GFP positive MCCs surrounded by DsRed positive outer cells indicating strong expression via the CMV promoter. In contrast, at ST48 we still find strong DsRed expression indicating CMV promoter activity, but we no longer see GFP positive cells (Figure 2B). While the embryos are transgenic for the lineage tracing construct, the Tub-CRE was injected as plasmid DNA, resulting in mosaic conversion. Consequently, not all MCCs are converted to green but all green cells are MCCs, as indicated by the presence of cilia with acetylated tubulin staining. Therefore, rather than quantifying the individual number of MCCs, we quantified the number of embryos in which we could identify any GFP + cells throughout developmental stages. At ST46 most embryos still have some GFP positive cells but by ST48 almost all embryos are devoid of GFP positive cells (Figure 2C). These results indicate that while some MCCs may transiently trans- or de-differentiate early, essentially all MCCs are lost from the epithelium by ST48. These results are consistent with our results using TgTub-memRFP (Figure 1B, 2C) which we utilize throughout the remainder of our experiments.

### Temporal shift from predominantly basal extrusion to an equal mix of apical and basal

During the period of time when MCC-driven fluid flow decreases between ST38 and ST48, we have observed two distinct forms of cell extrusion. First, as previously mentioned, we can observe the remnants of MCCs that have basally delaminated and undergone apoptosis, which remain visible for extended periods of time allowing us to quantify basal cell extrusion (Figure 3A, Movie S1). Additionally, we have observed a distinct pool of apically extruding live MCCs that are shed from the epithelium and undergo anoikis (Gudipaty and Rosenblatt, 2017; Stewart and Davis, 2019) (Figure 3B, Movie S2). Using long-term light sheet live imaging we have quantified the relative percentage of cells that undergo apical versus basal extrusion (e.g. Movie S3). Between ST42-46 we have observed that the majority of cells undergo basal extrusion and apoptosis (Figure 3D; ∼80% of extrusion events are basal). However, as development continues, we find a shift towards more apical extrusion, and after ST46 we find extrusion events are ∼50% apical and ∼50% basal (Figure 3D). Importantly, this shift towards apical extrusion in later embryos suggests the presence of distinct mechanisms driving apical versus basal extrusion of MCCs.

### Ectodermal caps show requirement for mesodermal signal in MCC loss

To investigate the mechanisms underlying MCC extrusion, we turned our attention to siganling pathways that govern MCC fate. Previously, Notch signaling has been shown to be a critical regulator of MCC loss (Tasca *et al*., 2021). Overriding normal signaling cues by driving the expression of the active Notch intracellular domain (NICD) specifically in MCCs using the Tub promoter leads to an early loss of MCCs via apoptosis. Additionally, blocking Notch signaling with the small molecule DAPT leads to an extended maintenance of MCCs. Importantly, the Notch ligand Jagged is expressed in the underlying mesoderm, suggesting that the relevant Notch signal emanates from below (Tasca *et al*., 2021). To further examine the role of Notch in MCC loss, we generated ectodermal “caps” by excising a portion of the ectoderm prior to the completion of gastrulation. This cap of cells balls up into a rough sphere that develops into a functional mucociliary organoid (Sokol and Melton, 1991; Walentek and Quigley, 2017; Weber et al., 1996). Importantly, in the absence of mesoderm, MCCs are maintained in these caps for a considerable time (Fukui et al., 2003). Strikingly, we find that at 15 dpf (ST50 of embryos) there are still a substantial number of MCCs (Figure 4A). Even after 1 month at ST55 we still find numerous MCCs although some TgTub-memRFP+ cells contain fewer or even no cilia present, suggesting that MCC maintenance wanes over time (Figure 4B). As expected, if we express NICD in MCCs using the Tub-promoter in caps we find that MCCs are again lost, further supporting the importance of Notch signaling and the underlying mesoderm in MCC fate regulation (Figure 4C-D) (Angerilli et al., 2018; Deblandre *et al*., 1999; Jones and Woodland, 1986; Stubbs et al., 2006). In this context, where embryo growth and elongation is not a contributing factor, these results suggests that Notch signaling is necessary for MCC loss.

**Figure 4:**
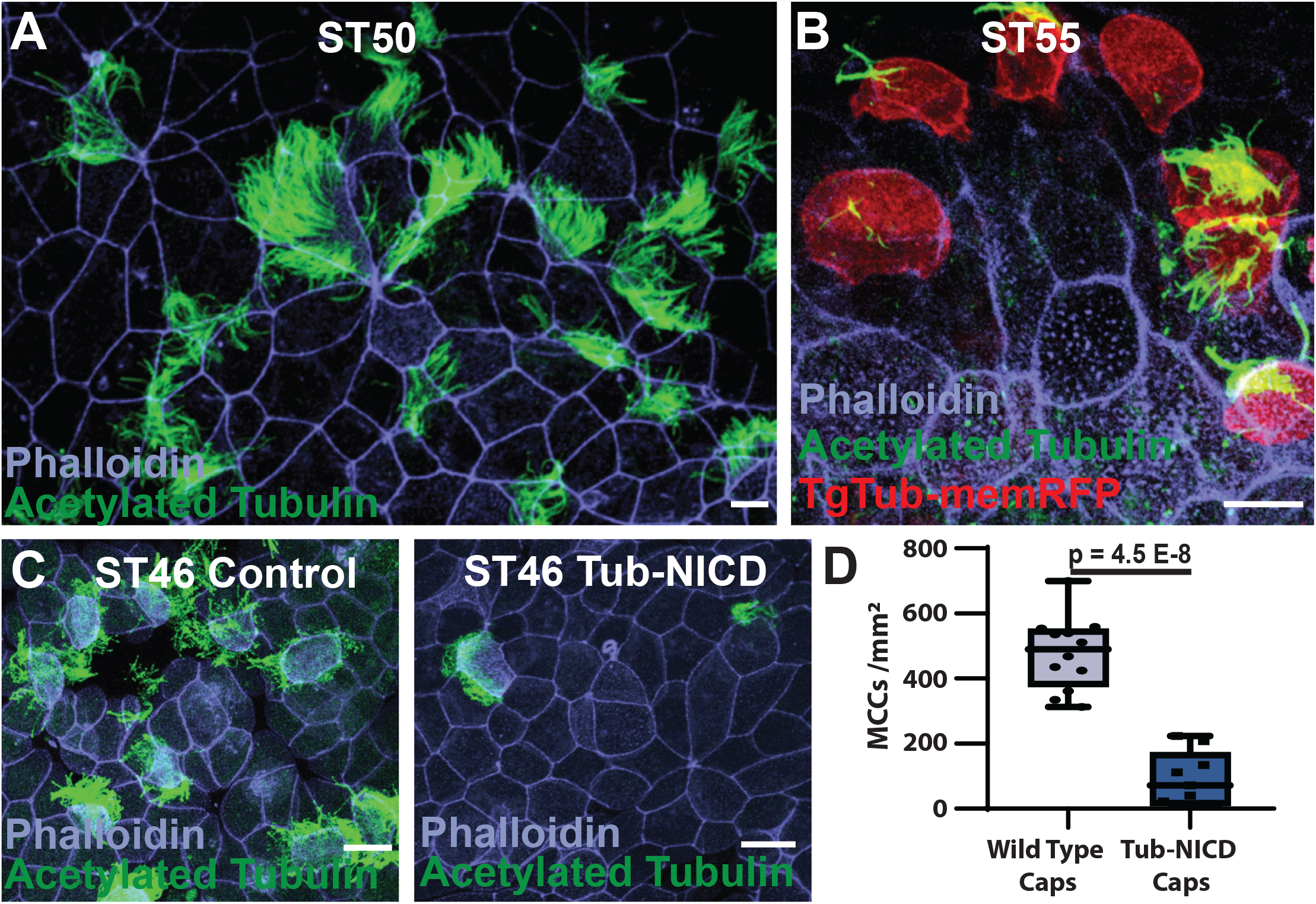
Mesodermal Notch is required for MCC loss. (A) Day 15 ST50 animal cap that still maintains a large number of MCCs. (B) Day 30 ST55 animal cap from a TgTub-memRFP embryo that shows the presence of multiple MCCs but also the loss of cilia and MCC maintenance in some RFP positive cells. (C) Comparison of MCCs in caps from WT embryos and embryos injected with Tub-NICD. (D) Quantification of MCCs in caps with and without Tub-NICD. Scale bar is 20μm.

### MCCs show age dependent receptivity to Notch

To determine if the underlying mesodermal signal is sufficient to drive MCC loss, we performed skin transplant experiments (Mitchell et al., 2009). We replaced small regions of skin from ST28 host embryos injected with mem-RFP with skin from ST11 donor embryos of a transgenic line expressing eGFP-OMP, a mitochondrial marker under control of the CMV promoter (Mitchell *et al*., 2009; Taguchi et al., 2012)(Figure 5A). We allowed the host embryos to develop to ST48, where most MCCs have been lost. Despite having the same underlying mesoderm, the younger donor tissue (now effectively ∼ST47) maintained many of its MCCs at a similar level to ST47 embryos and in contrast to the host tissue (ST48), which was largely devoid of MCCs (Figure 5B-C). Collectively, these results indicate that the mesoderm provides an instructive cue (via Notch ligands), but that there is a cell intrinsic, age-based feature that licenses the responsiveness to the Notch signal.

**Figure 5:**
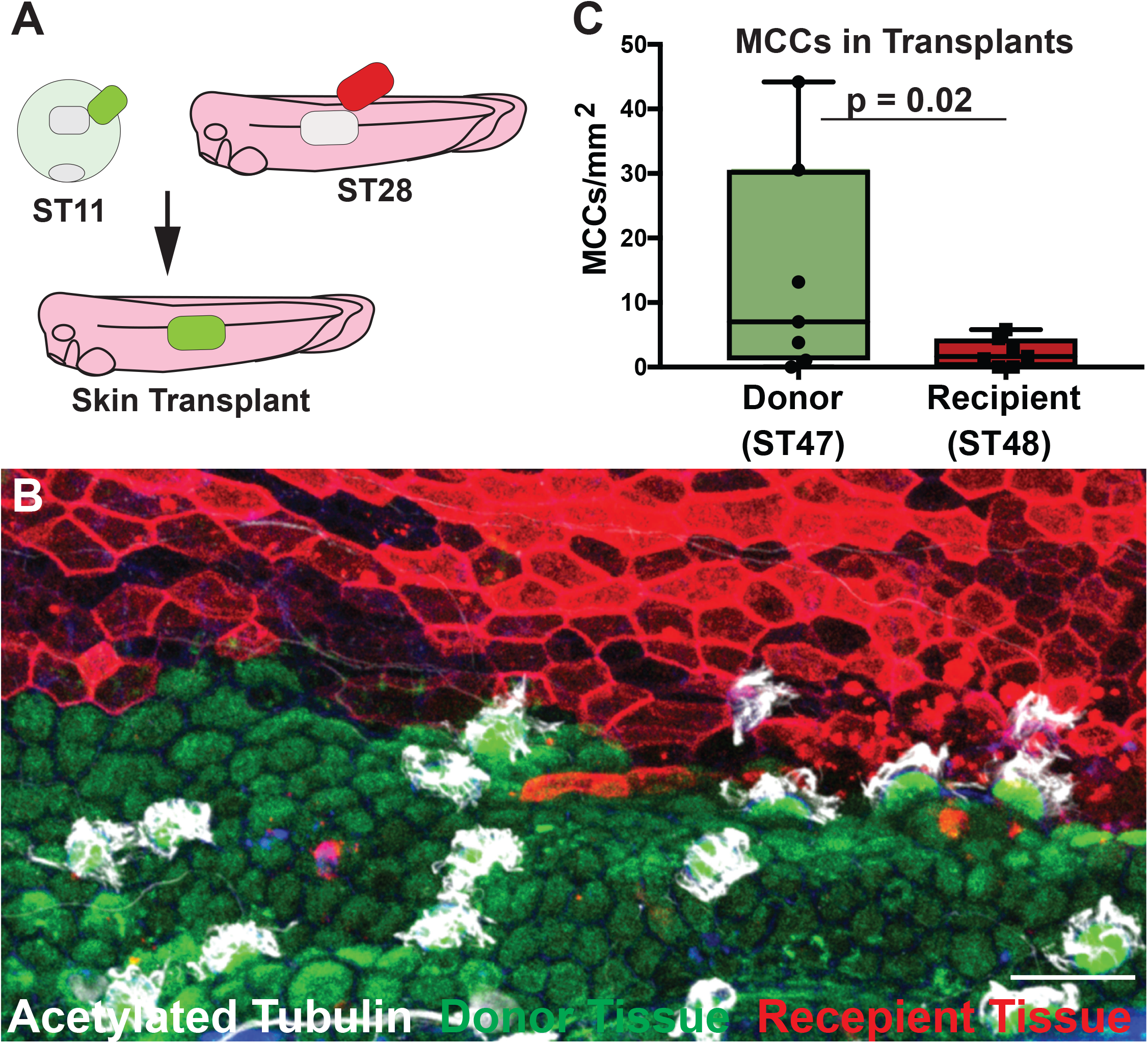
Skin transplants display age dependent loss of MCC. (A) Experimental design showing the transplantation of ST11 donor skin of *Xla*.*Tg(CMV:eGFP-OMP25)*^*Wtnbe*^ onto ST28 host skin of a CMV:memb-RFP injected embryo. (B) Representative image of an area containing both host tiusse (red, ST48) and donor tissue (green, ST47) stainied with acetylated tubulin (white). (C) Quantification of MCC number in host versus donor tissue, n=7 transplants, p=0.02. Scale bar is 50μm.

### MCC transcriptional program is protective against MCC loss

Our results show that the age of MCCs regulates their responsiveness to Notch, which led us to ask if we could manipulate the fate of MCCs by rebooting their transcriptional program. In the skin of *Xenopus* embryos, it is possible to deciliate MCCs using a brief treatment of high Ca^2+^ and low detergent (Werner and Mitchell, 2013). Importantly, MCCs will completely regrow their cilia within 4 HRs (Silva et al., 2016). This regrowth is blocked by the protein synthesis inhibitor cycloheximide, indicating that new transcriptional activity must occur to drive reciliation (Figure S1). We reasoned that if the ciliogenic program were restarted this may reset the age-based cell intrinsic Notch responsiveness. To test this, we deciliated embryos at ST28 and compared MCC number at ST48 with control non-deciliated animals. One might expect that this harsh treatment would make MCCs less healthy and more likely to undergo apoptosis. However, in contrast to controls, the deciliated embryos have a significant albeit modest maintenance of MCCs at ST48 (Figure 6A-C). Importantly, if we do this same treatment at ST46 right before the majority of MCC loss we see a more robust maintenance of MCCs indicating a temporal link between boosting the MCC transcriptional program and maintaining MCC lifespan (Figure 6D). These results suggests that in order for MCCs to undergo extrusion via the Notch pathway they must first abandon their MCC transcriptional program. This result is consistent with the transient trans- or de-differentation model that MCCs are losing their MCC fate prior to being extruded.

**Figure 6:**
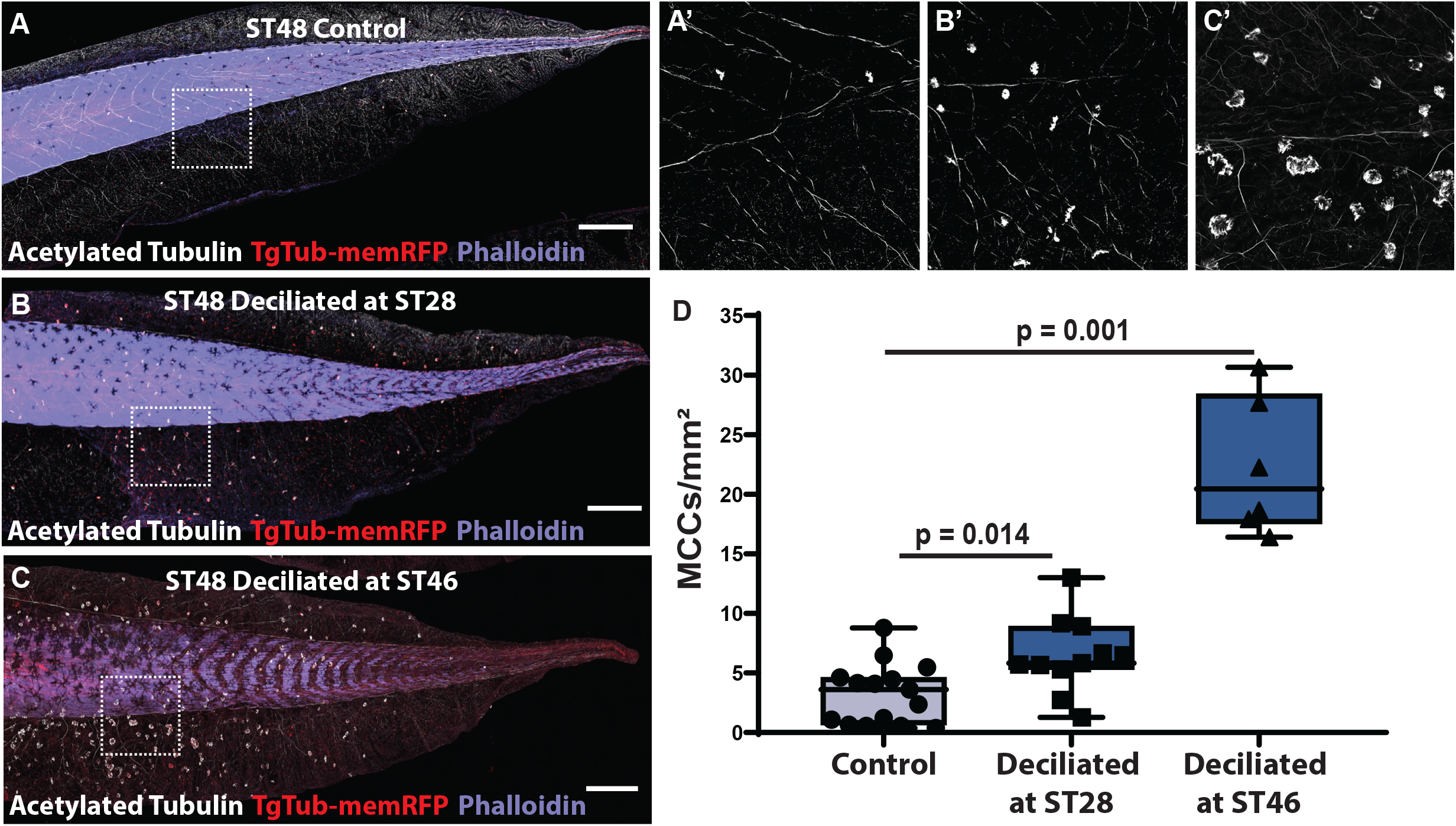
Deciliation of MCCs reboots transcriptional program and extends MCC lifespan. (A-C) Representative whole tail images of TgTub-memRFP (red) embryos stained with acetylated tubulin (white) and phalloidin (purple), showing MCCs in ST48 control embryo (A, A’), as well as ST48 embryos deciliated at ST28 (B,B’) and ST46 (C,C’). (D) Quantification of MCC number at ST48 in control and embryos that were deciliated at ST28 (n ≥ 8 embryos, p=0.014) or ST46 (n <u>=</u> 5 embryos, p = 0.001). Scale bar is 500μm. See Figure S1.

The transcriptional program driving MCC differentiation and ciliogenesis is well established (Collins et al., 2020; Lewis and Stracker, 2021; Walentek, 2021). In *Xenopus* embryos and vertebrate cell culture, the geminin family member MCIDAS has been shown to be essential for driving the formation of MCCs (Kim *et al*., 2018; Stubbs *et al*., 2012; Terre *et al*., 2016). Ectopic expression of MCIDAS in the ectoderm of *Xenopus* embryos converts all cells into MCCs and the depletion of MCIDAS leads to a loss of MCC formation (Stubbs *et al*., 2012). MCIDAS expression in *Xenopus* starts prior to MCC differentiation at ST12 but is largely turned off by ST26 suggesting that it is not essential for the short-term maintenance of the MCC fate (Briggs *et al*., 2018; Stubbs *et al*., 2012). However, given the important role of MCIDAS in initiating MCC fate we reasoned that extending its expression beyond ST26 could have profound effects on MCC fate maintenance and ultimately lifespan. We generated F0 transgenic embryos using a construct that drives expression of MCIDAS via the Tub promoter (Tub-MCIDAS). The Tub promoter is activated downstream of MCIDAS (Figgure S2), and therefore should generate a positive feedback loop maintaining MCIDAS expression. Similar to our deciliation experiments, the expression of MCIDAS provides a robust and significant maintenance of the MCC fate at ST48 (Figure 7). These results indicate that the maintenance of the ciliogenic program provides a protective feature against the loss of MCCs. We propose that the decommitment to the MCC fate is a licensing event that is required prior to cells responding to the Notch signal which drives basal extrusion and apoptosis.

**Figure 7:**
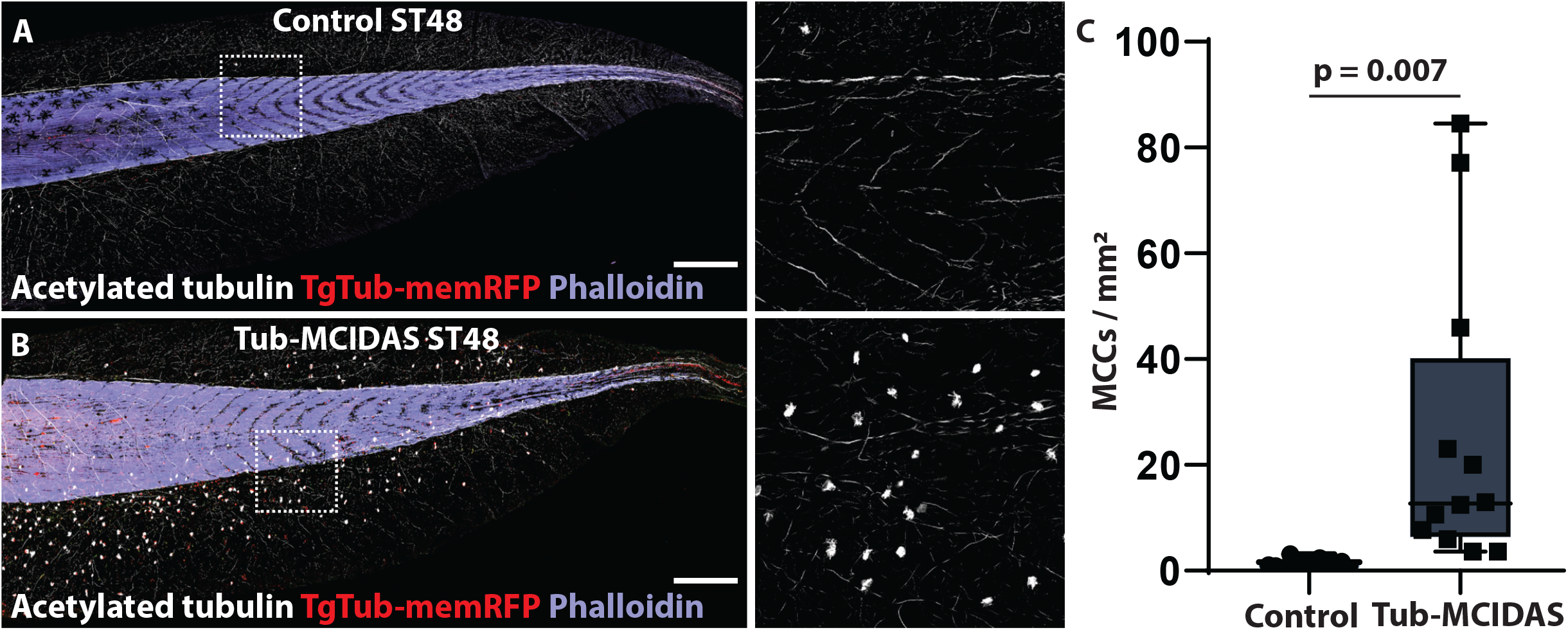
Maintenance of MCC fate protects against MCC loss. (A-B) Whole tail images of control and Tub-MCIDAS transgenic embryos with TgTub-memRFP (red), Acetylated tubulin (white) and phalloidin (purple). (C) Quantification of MCC number in control and Tub-MCIDAS embryos at ST48, n ≥ 12 embryos, p=0.007. Scale bar is 500μm. See Figure S2.

### Piezo 1 regulation of apical extrusion

The shift in preference towards apical extrusion in later embryos could represent a distinct mechanism for eliminating the remaining MCCs that have evaded Notch driven apoptosis. In some systems of cell extrusion, crowding rather than apoptotic signaling can be an initiator of cell loss (Eisenhoffer *et al*., 2012; Eisenhoffer and Rosenblatt, 2013; Franco et al., 2019). In this context, the mechanosensory channel Peizo 1 has been found to be an important regulator of extrusion. We reasoned that as development continues, cell crowding and the overall tissue remodeling associated with rapid growth between ST46-ST48 could be acting as a driver for the second wave of apical MCC extrusion (Zahn et al., 2022). To test this we first treated embryos with the Peizo 1 agonist Yoda1. At ST46, when DMSO treated embryos still maintain a considerable number of MCCs, we find that embryos treated with 100μM Yoda1 have significantly fewer MCCs (Figure 8A-C) (Botello-Smith et al., 2019; Lacroix et al., 2018; Syeda et al., 2015). Importantly, we do not find a significant difference in the number of RFP+ apoptotic vesicles indicating that the early loss of these MCCs was not at the expense of basally delaminating apoptotic cells (Figure 8D). This result suggests that activation of Piezo 1 is suffecient to drive apical extrusion.

**Figure 8:**
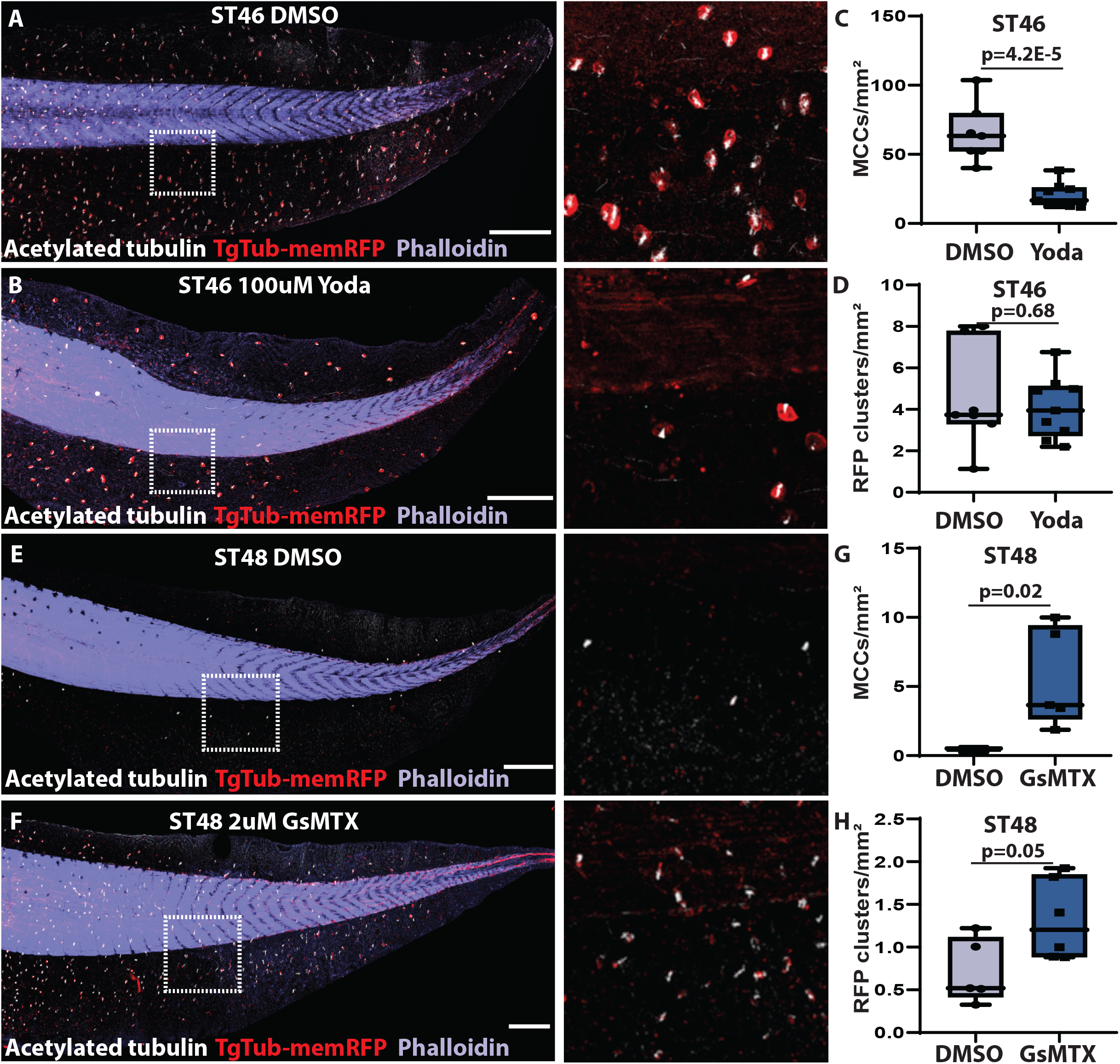
Piezo1 mechanosensation regulates MCC apical extrusion. (A-B) Representative images of DMSO treated (A) and 100uM Yoda treated (B) embryos with TgTub-mem-RFP (red), acetylated tub (white) and phalloidin (purple). (C) Quantification of MCCs (scored with acetylated Tub) in embryos treated with DMSO and 100uM Yoda, n ≥ 7 embryos, p=4.2E-5. (D) Quantification of RFP positive vesicles reflecting apoptotic MCCs in embryos treated with DMSO and 100uM Yoda, n ≥ 7 embryos, p=0.68. (E-F) Representative images of DMSO treated (E) and 2uM GsMTX treated (F) embryos with TgTub-mem-RFP (red), acetylated tub (white) and phalloidin (purple). (G) Quantification of MCCs (scored with acetylated Tub) in embryos treated with DMSO and 2uM GsMTX, n ≥ 8 embryos, p=0.02. (H) Quantification of RFP positive vesicles reflecting apoptotic MCCs in embryos treated with DMSO and 2uM GsMTX, n ≥ 5 embryos, p=0.05. Scale bar is 500μm.

We next tested the Peizo 1 inhibitor GsMTX and found that at ST48 when DMSO treated embryos were largely devoid of MCCs there is a significant maintenance of MCCs in embryos treated with 2μM GsMTX suggesting Piezo 1 is also necessary for apical extrusion (Figure 8E-G)(Bae et al., 2011). In this context we do find a slight increase in RFP+ vesicles, that suggests that these populations are at least partially overlapping (Figure 8H). While driving early mechanosensory apical loss would not be expected to affect Notch driven basal loss, the reverse is not as likely. Given that these modes of extrusion overlap, we would expect that blocking mechanosensory loss via GsMTX would lead to more cells that are available to be responsive to Notch later and therefore driven towards basal extrusion. Overall, these results suggest that there are two independent mechanisms for MCC extrusion, an apoptotic basal extrusion driven by Notch signaling and a live apical extrusion driven by a Peizo 1 mechanosensory mechanism.

## Discussion

With each MCC containing ∼150 cilia that each beat at ∼15 Hz, the epithelial fluid flow that MCCs generate is extremely energetically taxing to the embryo. When flow is no longer needed due to the swimming motion of the tadpole, there would presumably be a strong selective pressure to remove these cells. Notch signaling is a critical regulator of ciliated epithelia both during development as well as during tissue remodeling after insult (Deblandre *et al*., 1999; Lafkas *et al*., 2015; Liu *et al*., 2007; Rock *et al*., 2011; Tasca *et al*., 2021). During *Xenopus* MCC loss, Notch is a critical signal that pushes MCCs towards apoptosis and basal extrusion. Here we show that the receptivity to the Notch signal is regulated by the overall MCC transcriptional state of the cell. This is based on three experiments. First, skin transplants of younger skin do not respond with the same kinetics as older host tissue. Second, deciliated MCCs reboot their MCC transcriptional program and this offers resistance to the Notch signal, extending the MCC lifespan. Finally, continued expression of the MCC transcriptional regulator MCIDAS, maintains the MCC transcriptional program and extends MCC lifespan. Our data suggests a balance between MCC fate maintenance and the ability to respond to the Notch signal. We propose a licensing mechanism where MCCs must first decommit from their fate prior to becoming responsive to Notch driven apoptosis. In all of our experiments the Notch ligand should be consistently present, suggesting that the change as MCCs decommit is likely due to either a change in Notch receptor expression level or Notch receptor subtype specificity and sensitivity. This is consistent with our animal cap data as well as previously published work that the expression of the constitutively active NICD in MCCs is sufficient to drive apoptosis (Tasca *et al*., 2021). Additionally, miR-449 is part of the MCC transcriptional program and is known to negatively regulate the Notch pathway which could explain why decommiting from the MCC fate leads to a change in Notch sensitivity (Marcet NCB2011).

Cell extrusion is an important mode of tissue remodeling that is critical for removing unwanted, less competitive, or dying cells (Eisenhoffer and Rosenblatt, 2013; Gudipaty *et al*., 2018; Mitchell and Rosenblatt, 2021). Cell extrusion has been widely studied and there are an extensive number of tissues that have been shown to exhibit cell extrusion. What makes the extrusion of MCCs somewhat distinct is that a specific cell type that is interspersed across the epithelium is lost within a defined developmental period by two distinct yet overlapping mechanisms. Even more unusual is that the direction of extrusion is different between these two mechanisms. In most systems cell extrusion is normally unidirectional and only occurs bidirectionally in disease or experimental conditions. Yet MCCs naturally undergo both Notch driven basal extrusion as well as Piezo 1 regulated apical extrusion during an overlapping developmental window. We propose that MCCs will respond to the Notch signal and apoptose if they have decommitted from their MCC fate. If they do not respond, due to differential age (e.g. some MCCs are born later than others) or due to potential unfavorable positions relative to the source of the Notch ligand, they will be driven to apical extrusion via mechanosensation. Mechanosensory-driven cell extrusion has been described in a variety of tissues, yet the extruding cells are typically thought to be in an energetically unfavorable position (e.g. the edge of the tissue). MCCs represent cells with a significant cytoskeletal architecture at their apical surface that could provide a distinct environment which facilitates apical extrusion. Another possibility is that as MCCs decommit from their fate they become less energetically distinct from their neighbors and represent a sickly, less competitive cell susceptible to extrusion. Alternatively, these remaining MCCs could maintain their fate, maintain their cytoskeletal architecture, and as such be targeted for extrusion based on their energetically distinct cellular profile. Distinguishing between these alternatives will be an important aspect of future work.

## Methods

### Transgenic Xenopus and Embryo Injections

All *Xenopus* experiments were performed using previously described techniques (Werner and Mitchell, 2013). *In vitro* fertilizations were performed using standard protocols (Sive et al., 1998; 2007b; 2010) that have been approved by the Northwestern University Institutional Animal Care and Use Committee. Transgenic *Xenopus* that express membrane-bound RFP driven by the tubulin promoter (Xla.Tg(tuba1a:MyrPalm-mRFP)^NXR^), were previously generated and obtained from the National *Xenopus* Resource Center (NXR) (Brooks and Wallingford, 2015). For skin transplants, transgenic *Xenopus* that express eGFP tagged OMP driven by the CMV promoter (Xla.Tg(CMV:eGFP-OMP25)^Wtnbe^)allowing for constitutive GFP expression in mitochondrial membranes were obtained from NXR (Taguchi *et al*., 2012). Wild type or transgenic embryos were injected at the two- or four-cell stage with 40-250 pg mRNA or 10-20 pg of plasmid DNA.

### Plasmids and mRNA

In this study, we employed a CMV promoter to drive constitutive expression of GFP or RFP as tracers in all cell types, while MCC specific expression was driven by an alpha tubulin (aTub) promoter, as previously described (Stubbs *et al*., 2006).

I-sceI-CMV-LoxP-DsRed-LoxP-GFP: To generate the I-sce-CMV-DsRed-LoxP-GFP cassette, CMV-LoxP-DsRed-LoxP-GFP was amplified from pLV-CMV-LoxP-DsRed-LoxP-GFP (Addgene #65726) (Zomer et al., 2015) and inserted into an I-sceI vector, a gift from Marko Horb at Marine Biological Laboratory (MBL), by Gibson Assembly.

pCS2-aTub-Cre2 and I-sceI-Tub-Cre2: The CMV promoter of pCS.Cre2 (Addgene #31308) (Ryffel et al., 2003) was replaced with an alpha-tubulin promoter by SalI(5’)/HindIII(3’). To insert into the I-sceI vector, aTub-Cre2 was cut out and ligated into an I-sceI vector by SalI(5’)/KpnI(3’).

pCS2-aTub-NICD-GFP: Notch intracellular domain (NICD) (Deblandre *et al*., 1999) was inserted into the pCS2-aTub-GFP construct using ClaI(5’)/XbaI(3’).

I-sce-aTub-GFP-MCIDAS: The CMV promoter of pCS2-MCIDAS (Stubbs et al., 2012) was replaced with aTub-GFP by SalI(5’)/EcoRI(3’) to generate pCS2-tub-GFP-MCIDAS. Then, aTub-GFP-MCIDAS was inserted into the I-sceI vector using the SalI(5’)/KpnI(3’) sites.

pCS2-memb-RFP (Stubbs *et al*., 2006) was used for skin transplants, and mRNA of LifeAct-GFP (Werner and Mitchell, 2013) was synthesized with the Sp6 mMessage Machine kit (Life Technologies, AM1340) and purified by RNeasy MiniElute Cleanup Kit (QIAGEN, 74204).

### I-sceI transgenesis

To generate transgenic embryos, we utilized I-sceI meganuclease-mediated method as previously described (Ishibashi et al., 2012; Ogino and Ochi, 2009; Pan et al., 2006). I-sceI-CMV-LoxP-DsRed-LoxP-GFP construct was incubated with I-sceI at 37’C for 40min and embryos were injected with 40pg of DNA and 0.004U of I-sceI per embryo at the one-cell stage until the 1hour mark immediately followed by dejelling. To optimize transgenesis efficiency and minimize toxicities from a high amount of DNA and enzyme injections, we used only one I-sceI construct for the CMV-LoxP-DsRed-LoxP-GFP cassette (Thermes et al., 2002). Then, embryos were allowed to heal and pCS2-aTub-Cre was injected at the 4-cell stage. Following transgenesis at ST38, the Leica M165 FC dissecting microscope was used to isolate embryos for lineage tracing that were positive for GFP expression, indicating DsRed to GFP conversion in MCCs. The percentage of these embryos that remained GFP-positive were scored at various developmental stages.

### Fluid flow measurement

Fluorescent microspheres were used to visualize cilia-driven fluid flow at different stages of development as previously described using the Leica M165 FC dissecting microscope connected to the Casio Exilim 60fps mounted digital camera (Werner and Mitchell, 2013). In brief, fluorescent microspheres (#F8836, Invitrogen Inc.) were resuspended in 0.1X MMR containing 0.5% glycerol. Embryos were placed in 0.02% tricaine+0.1X MMR and fluorescent beads were added dropwise to the embryo. The relative velocity of the fluid flow was calculated by determining the displacement of individual microspheres over the surface of the embryo by using simple particle tracking software available on FIJI (Schindelin et al., 2012).

### Immunostaining

Embryos were fixed with 3% PFA/PBS and blocked in 10% heat-inactivated goal serum (HIGS)/PBS after washing with PBS. To visualize cilia, embryos were incubated with mouse anti-acetylated α-tubulin (T7451; Sigma-Aldrich, 1:500) in 5% HIGS/PBS for an hour at room temperature. Then, Cy-2- or Cy-5-conjugated goat anti-mouse secondary antibodies (Thermo Fisher Scientific) were used at a 1:750 dilution in 5% HIGS/PBS. To visualize actin and nuceli, Phalloidin 650 (PI21838, Invitrogen, 1:300) and DAPI (#62248, Thermo Fisher Scientific, 1:300) were used. Following staining, embryo tails were mounted between two coverslips as previously described (Werner and Mitchell, 2013) using Fluoro-Gel (#1798510, Electron Microscopy Sciences).

### Microscopy

Fixed samples were imaged using the Nikon A1R laser scanning confocal microscope using a 60x oil Plan-Apochromat objective lens with a 1.4 NA. For whole tail imaging, a 20x water Plan Flour objective lens with a 0.75 NA with Large Image function was utilized. Maximum intensity projections of z-stacks were analyzed using either Nikon NIS Elements or FIJI (Schindelin *et al*., 2012). Live imaging of apical and basal extrusion was performed using the Nikon A1R laser scanning confocal microscope with a 60x oil Plan-Apochromat objective lens with a 1.4 NA. Long term imaging for quantification was performed on a Nikon Ti2 micropscope equipped with a Mizar TILT light sheet and a photometric prime 95B camera, using a 10X objective (N.A. 0.3) or 100X silicone objective (N.A. 1.35). Images were analyzed using FIJI software for the calculation of the ratio between apical and basal extrusion.

### Ectodermal caps

Embryonic ectodermal caps were isolated at ST10 in 0.5X MMR+gentamicin, following a previously described protocol (Sive et al., 2007a). Caps were then transferred to fresh 0.5x MMR + gentamicin to recover. After healing, caps were treated with 0.5ng/mL of Activin A (A4941-10, Sigma-Aldrich) in 0.5x MMR + gentamicin for one hour to promote development to epidermis (Ariizumi et al., 1991). Caps were transferred to Leibovitz L-15 cell culture medium+ Steinberg+1%BSA+antibiotics for long term culture (Ariizumi et al., 2017; Fukui *et al*., 2003). Uncapped embryos were used as a stage reference. Once caps are grown for two weeks (∼ST50) and a month (∼ST55), caps were fixed in 3% PFA/PBS, followed by antibody staining.

### Skin transplants

Embryonic skin transplants were performed as previously described (Mitchell *et al*., 2009; Werner and Mitchell, 2013). Briefly, a small region of ectoderm from a donor embryo (Xla.Tg(CMV:eGFP-OMP25)^Wtnbe^) at ST10 was removed with a fine hair then transplanted onto the recipient embryo at ST28 injected with pCS2-memb-RFP after removing a similar patch of outer skin layer. All transplant experiments were performed in Danilchik’s buffer + 0.1% BSA + 0.02% Tricaine. Then, transplanted tissue was held in place by gently pressing down with a small piece of glass coverslip using silicone grease to secure its position for approximately 1∼2 hours. After healing, the coverslip was gently removed, and embryos were grown in 0.1X MMR until the desired stage was reached.

### Deciliation

Following a method previously described (Werner and Mitchell, 2013), embryos were deciliated at various developmental stages in 0.1x MMR + gentamicin containing 75 mM calcium and 0.02% NP40. Embryos were incubated in this solution for approximately 20 - 40 seconds and then washed several times in 0.1x MMR + gentamicin. Several control embryos were immediately fixed in 3% PFA/PBS following treatment to confirm successful deciliation. The remaining embryos were incubated in 0.1x MMR + gentamicin until reaching the desired stage, then fixed in 3% PFA/PBS for analysis.

### Drug treatments

Yoda1 (Fisher, #558610) was used to activate Piezo1 mechanosensory channels, whereas GsMTX (MedChemExpress, #HY-P1410) was used to inhibit Piezo1 channels. Embryos were incubated in the presence of DMSO (vehicle), 100 μM Yoda (in DMSO), or 2 μM GxMTX (in DMSO) beginning at ST28 of development until the embryos reached the desired stage. Then, embryos were fixed in 3% PFA/PBS.

## Acknowledgments

This work was supported by NIH/NIGMS to B.J.M. (R01GM089970). R.V. was supported from the Cutaneous Biology training grant (TGM32AR060710). O.H. was supported off a diversity supplement from (R01GM119322). P.W. was supported by the Deutsche Forschungsgemeinschaft (DFG) under the Emmy Noether Programme (WA3365/2-1) and under Germany’s Excellence Strategy (CIBSS–EXC-2189–Project ID 390939984). We would like to thank the National *Xenopus* Resource and the Marine Biological Laboratories for technical support and reagents.

## Data Availability

There are no large datasets associated with this work, but any imaging data is available upon request.

## Figure Legends

**Figure S1:**
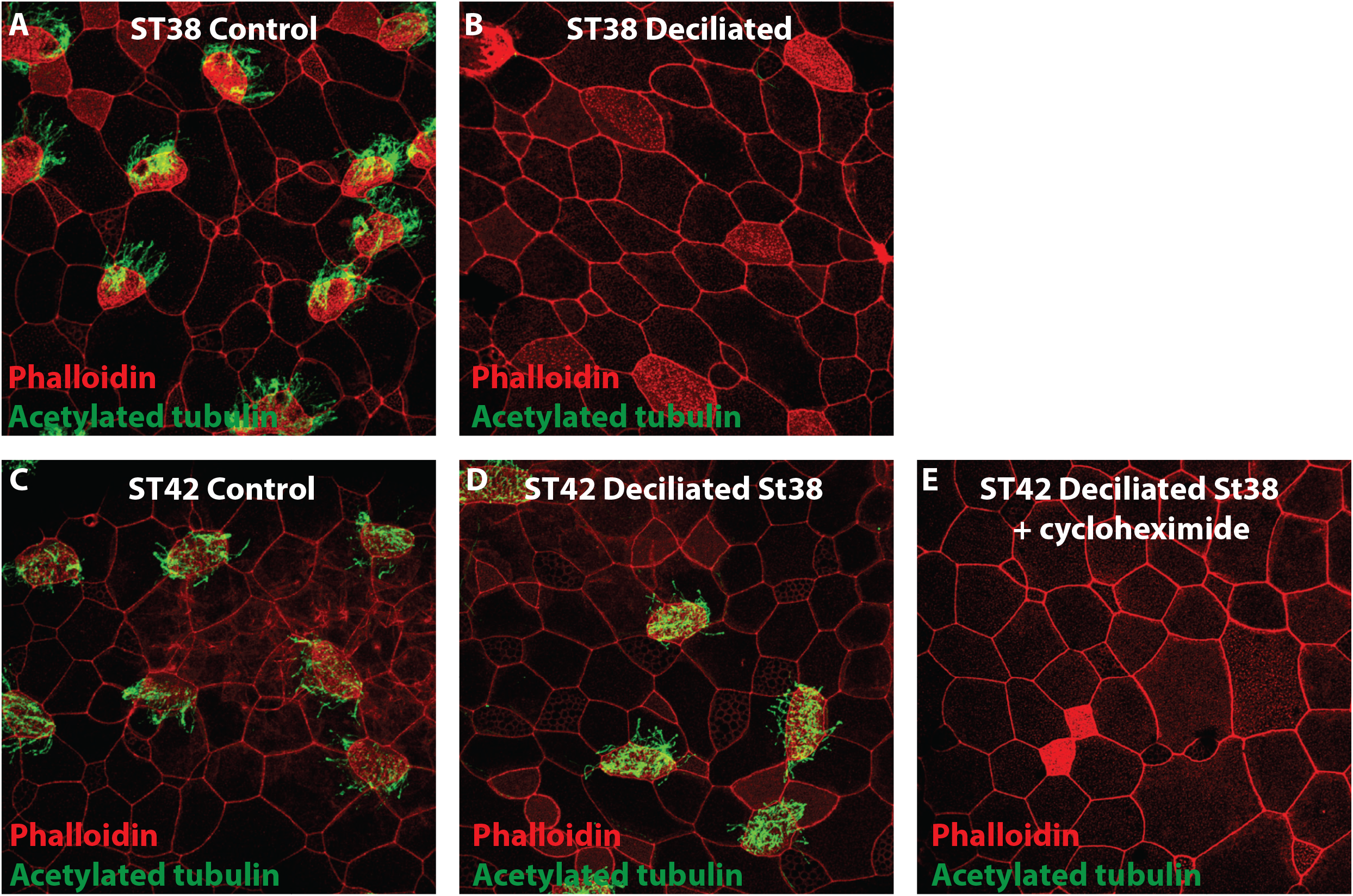
Multiciliated cell deciliation. (A-E) Representative images of a embryos stained with phalloidin (red) and acetylated tubulin (green). (A-B) Representative images of a control embryo at ST38 (A) and a ST38 embryo directly after deciliation (B). (C-E) Representative images of a control embryo at ST42 (C), a ST42 embryo that was deciliated at ST38 (D), and an embryo that was decilated at ST38 but that was also treated with cycloheximide (E).

**Figure S2:**
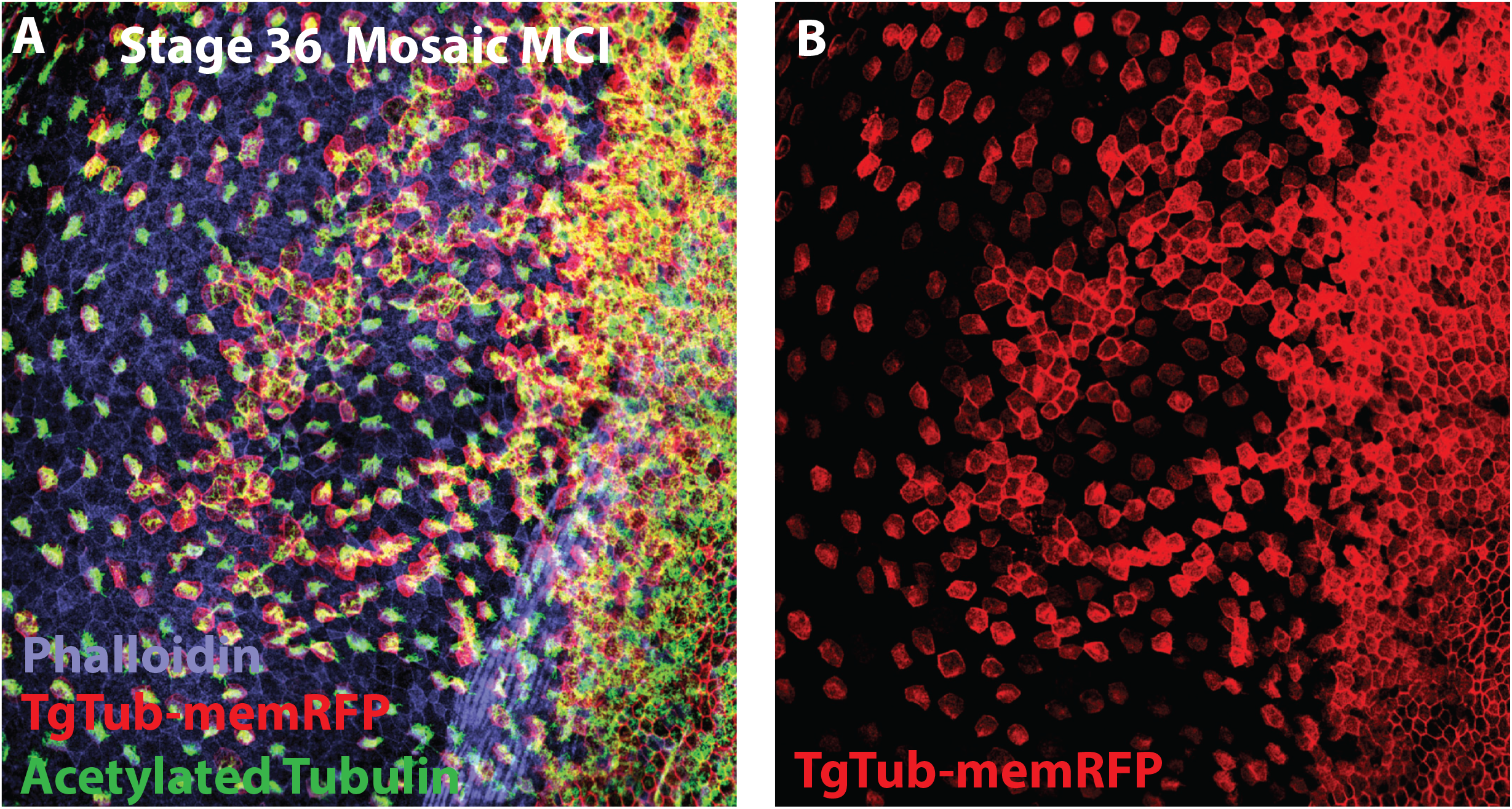
Tub promoter activity is downstream of *Mcidas*. (A-B) ST36 embryo mosaically injected with the MCC inducing factor Mcidas, showing broad expression of the TgTub-memRFP (red) in extopic MCCs labled with acetylated tubulin (green) with phalloidin staining (purple) (A) and the same image showing only the ectopic TgTub driven mem-RFP (B).

**MovieS1: MCC basal extrusion**. This is a top down projection of the data in Figuire 3A of an embryo expressing TgTub-memRFP and LifeACT-GFP showing a cell basally extruding. Movie is 35 frames with images taken every 20 minutes.

**MovieS2: MCC apical extrusion**. This is a top down projection of the data in Figuire 3B of an embryo expressing TgTub-memRFP and LifeACT-GFP showing a cell apically extruding. Movie is 25 frames with images taken every 10 minutes.

**MovieS3: MCC apical and basal extrusion**. Cropped image from a low resolution (10X) long-term light-sheet movie that shows how we quantified apical extrusion (arrow frame 15) and basal extrusion (arrowhead frame 81). Movie is 100 frames with images taken every 3 minutes.

